# Real-time alerts from AI-enabled camera traps using the Iridium satellite network: a case-study in Gabon, Central Africa

**DOI:** 10.1101/2021.11.10.468078

**Authors:** Robin C. Whytock, Thijs Suijten, Tim van Deursen, Jędrzej Świeżewski, Hervé Mermiaghe, Nazaire Madamba, Narcys Moukoumou, Joeri A. Zwerts, Aurélie Flore Koumba Pambo, Laila Bahaa-el-din, Stephanie Brittain, Anabelle W. Cardoso, Philipp Henschel, David Lehmann, Brice Roxan Momboua, Loïc Makaga, Christopher Orbell, Lee J.T. White, Donald Midoko Iponga, Katharine A. Abernethy

**Affiliations:** Faculty of Natural Sciences, University of Stirling, FK9 4LA, UK; Hack the Planet, Q42, The Netherlands; Appsilon AI for Good, Warsaw, Poland; School of Architecture and Environment, Department of Landscape Architecture University of Oregon, Eugene, OR 97403-5249 US; Agence Nationale des Parcs Nationaux, Libreville, Gabon; Utrecht University, Heidelberglaan 8, 3584 CS Utrecht, The Netherlands; School of Life Sciences, University of KwaZulu-Natal, South Africa; University of Oxford, Department of Zoology, Oxford, UK; Department of Ecology and Evolutionary Biology, Yale University, New Haven, CT 06520, USA; Panthera, 8 West 40th Street, 18th Floor, New York, NY 10018, USA; Ministry of Water and Forests, Boulevard Triomphal, Libreville, Gabon; Institut de Recherche en Ecologie Tropicale, CENAREST, BP 842 Libreville, Gabon

## Abstract

1. Efforts to preserve, protect, and restore ecosystems are hindered by long delays between data collection and analysis. Threats to ecosystems can go undetected for years or decades as a result. Real-time data can help solve this issue but significant technical barriers exist. For example, automated camera traps are widely used for ecosystem monitoring but it is challenging to transmit images for real-time analysis where there is no reliable cellular or WiFi connectivity. Here, we present our design for a camera trap with integrated artificial intelligence that can send real-time information from anywhere in the world to end-users.
2. We modified an off-the-shelf camera trap (Bushnell™) and customised existing open-source hardware to rapidly create a ‘smart’ camera trap system. Images captured by the camera trap are instantly labelled by an artificial intelligence model and an ‘alert’ containing the image label and other metadata is then delivered to the end-user within minutes over the Iridium satellite network. We present results from testing in the Netherlands, Europe, and from a pilot test in a closed-canopy forest in Gabon, Central Africa.
3. Results show the system can operate for a minimum of three months without intervention when capturing a median of 17.23 images per day. The median time-difference between image capture and receiving an alert was 7.35 minutes. We show that simple approaches such as excluding ‘uncertain’ labels and labelling consecutive series of images with the most frequent class (vote counting) can be used to improve accuracy and interpretation of alerts.
4. We anticipate significant developments in this field over the next five years and hope that the solutions presented here, and the lessons learned, can be used to inform future advances. New artificial intelligence models and the addition of other sensors such as microphones will expand the system’s potential for other, real-time use cases. Potential applications include, but are not limited to, wildlife tourism, real-time biodiversity monitoring, wild resource management and detecting illegal human activities in protected areas.

## Introduction

Goals towards biodiversity protection, the sustainable use of ecosystems, and mitigation of climate change are now clearly defined for nearly every nation on earth (Convention on Biological Diversity, 2021; UN General Assembly, 2015). However, efforts to protect and preserve ecosystems are often hindered by long delays (months, years or more) between the timing of data collection and data analysis. Ecosystem change and ecosystem threats can therefore go undetected for extended periods. Affordable technology for real-time ecosystem monitoring and threat detection could help address this issue, but significant technological barriers exist. In particular, it has proven a challenge to generate reliable, real-time data from some sensors such as automated camera traps in the absence of wireless fidelity networks (WiFi) or broadband cellular networks.

Automated camera traps (or ‘trail cameras’) are used to detect and survey wildlife and by conservation managers to identify ecosystem threats (Bessone et al., 2020; Hobbs & Brehme, 2017; Wearn & Glover-Kapfer, 2019). A typical camera trap comprises a movement or heat sensor (e.g. a passive infra-red sensor), one or more digital image sensors, a flash or night-vision capability, removable digital storage and a battery power source. Many commercial models are available and cameras can also be easily custom-built using off-the-shelf components (Droissart et al., 2021).

Network-enabled camera traps, which send captured images to users in real-time, are now commercially available but typically need access to a reliable broadband cellular network connection. In many countries, however, cellular network coverage is still limited and is often unreliable, causing ‘data poverty’ (Leidig & Teeuw, 2015). Cellular network coverage is also usually focused on human population centres, which might be far from areas of ecological or conservation interest. As a result, camera traps with network connectivity are rarely deployed at scale in these network-limited landscapes.

In network-limited landscapes, there have been some attempts to use WiFi or GSM enabled camera traps by building dedicated infrastructure such as communication towers and meshed networks. These systems transmit the images over the network for later analysis. However, it can be prohibitively expensive to build the necessary infrastructure and it is often logistically impossible in the most rugged landscapes. Legal barriers also exist and commercial providers can own the exclusive rights to build and install GSM towers and transmitters. Satellite networks have the best global coverage, but high data transfer costs mean it is expensive to send images generated by camera traps to end-users in real time.

Beyond network connectivity, another challenge limiting the usefulness of camera traps for timely decision-making has been extracting relevant information from the image, or “image labeling”. In ecology, images are typically labelled by identifying the species in the image and counting the number of individuals seen. Camera trap projects collect large volumes of data and it is not uncommon to generate millions of images or videos that require terabytes of storage space. Solutions to labeling these large image databases range from using dedicated software that speeds up manual image labeling, to large-scale citizen science projects and the use of artificial intelligence algorithms (Beery et al., 2019; Swanson et al., 2016). The precision and accuracy of the latest artificial intelligence algorithms for image labelling now approach or match human experts for some species but they typically require powerful computing resources either based on ‘the cloud’ or locally using expensive hardware (Norouzzadeh et al., 2018; Tabak et al., 2019; Whytock et al., 2021). However, recent developments in the field of ‘edge computing’ allow artificial intelligence algorithms to be deployed on microcomputers with relatively low computing and electrical power requirements. It is therefore possible to integrate artificial intelligence with camera trap hardware for deployment in the field. These advances mean that data-light image labels generated by artificial intelligence algorithms can be inexpensively transmitted over wireless networks (e.g. satellite) instead of the costly, data-heavy images.

Here, we present an overview of a ‘smart’ camera trap system that integrates artificial intelligence with a popular off-the-shelf camera trap for real-time alerts over the Iridium satellite network. The system also transmits information on power status, temperature and humidity for the purposes of monitoring hardware integrity. Although the system is based on existing (open source) hardware where possible, our aim is not to provide a blueprint for a finished ‘tool’, such as the Audiomoth bioacoustic recorder (Hill et al., 2018), but to provide insights into how we solved significant technical challenges.

Individual off-the-shelf components can also rapidly change or become unavailable (e.g. components for a bioacoustic recorder (Whytock & Christie, 2017)), potentially making it difficult for end-users to follow blue-print designs. As with all surveillance systems, including existing camera trap technology, there are significant ethical and legal issues to consider before using smart cameras in the field, particularly where human subjects may be intentionally or unintentionally observed (Sandbrook et al., 2018). We therefore caution that deployment of the technology presented here should be guided by robust ethical review.

To evaluate the system’s effectiveness, we present systematic results from testing in the Netherlands and a field test in a high-canopy tropical forest in Gabon, Central Africa. In Gabon, we deployed five systems for real-time detection of forest elephant *Loxodonta cyclotis* with the long-term aim of using the system to help mitigate forest elephant crop depredation incidents. These incidents are a pressing concern for the country’s success in aligning conservation objectives with rural development. Other uses for which the system could also be used, such as real-time wildlife monitoring and detecting illegal human activities such as poaching, are also discussed.

## Methods

### General summary

Our objective was to create a robust, field-ready system that could (1) provide real-time alerts from camera traps at an affordable cost, (2) be deployed in the most rural landscapes without existing GSM, Long Range radio (LoRa) or WiFi coverage, (3) function without installing additional infrastructure such as communication towers, base stations or meshed networks, (4) be easily deployed by users who do not have a specialist background in using artificial intelligence-enabled technology and (5) avoid re-inventing existing technology (e.g. camera traps), thus allowing us to solve the problem within a relatively short time frame.

Our solution was to modify a standard Bushnell™ camera trap by adding additional hardware allowing it to communicate wirelessly with separate, self-contained computing resources installed nearby - which we named the ‘smart bridge’ (Figure 1). The smart bridge is based on an earlier prototype designed to take photographs of wild penguins (https://github.com/IRNAS/arribada-pmp), and provides an intelligent link, or ‘bridge’, between the camera trap and the end user.

**Figure 1.**
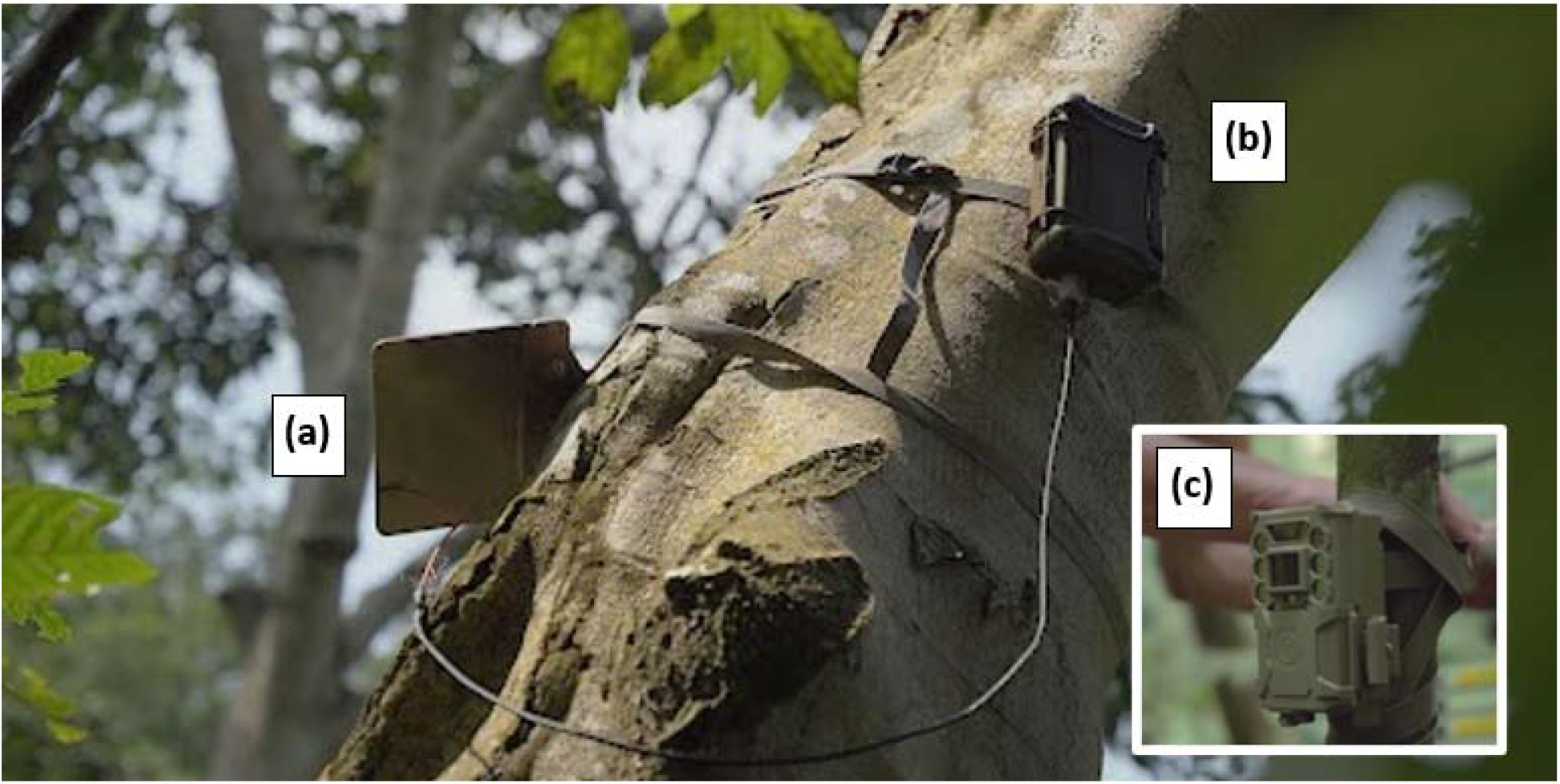
System deployed in the field showing the solar panel (a) and smart bridge (b) attached to a tree approximately 6 m above ground level. The Bushnell™ camera trap (c) is installed at ground level approximately 10 m away from the smart bridge.

We customised the camera trap by installing a microcontroller with LoRa capabilities based on the OpenCollar Lion Tracker (https://github.com/IRNAS/smartparks-lion-tracker-hardware). Instead of the standard secure digital (SD) card, we used a WiFi-enabled SD card. When an image is captured by the camera trap, the LoRa board in the camera alerts the smart bridge and activates the WiFi SD card, creating a local WiFi network. The smart bridge boots a Raspberry Pi Compute Module 4 that joins the WiFi network and retrieves the image or images from the camera. The species contained in the image are then identified using an artificial intelligence algorithm for species classification. The species and metadata associated with the image (time, date, location) and smart bridge sensor data (internal temperature, humidity and power status) are finally transmitted in an encoded message from the smart bridge to a web-based application running in the cloud (Google’s App Engine). The data are sent over the Iridium satellite network, which provides global coverage within minutes. To save power, the Raspberry Pi then shuts down and the smart bridge enters a low-power sleeping mode. Pairing between the camera and smart bridge is automatic and requires no user input or setup. A diagram of the system logic is shown in Figure 2.

**Figure 2.**
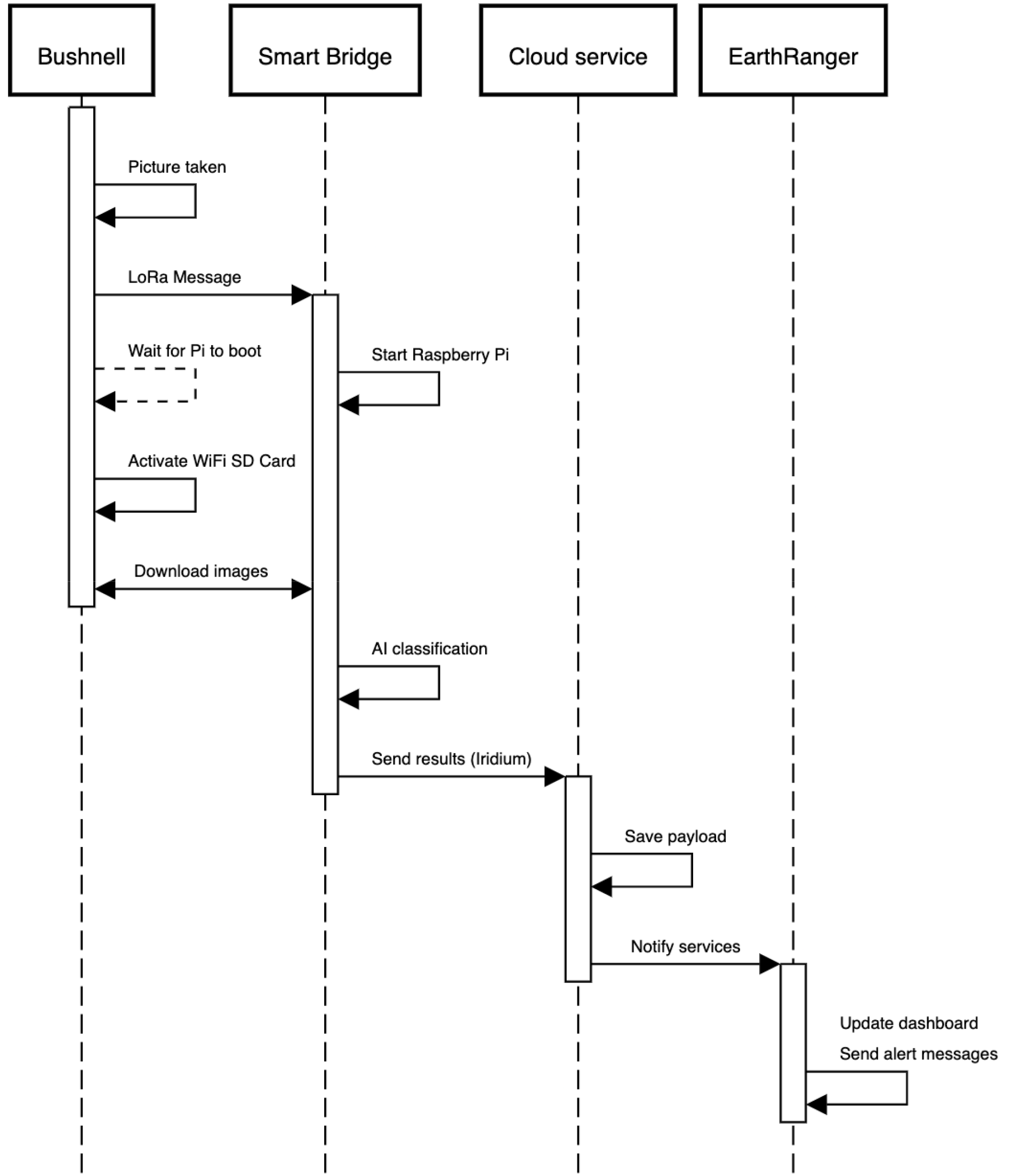
Diagram showing the stepwise logic between the Bushnell™ camera trap capturing an image and sending an alert via the smart bridge. Total duration of the entire process is approximately five minutes under optimal conditions.

## Hardware stack

### Camera

We used a Bushnell™ Core 24MP Low Glow 119936C camera trap for development but similar modifications can be made to other models and brands. The camera was set to take single images (2304 × 1296 pixels, 72 dpi) at 10s intervals with sensitivity set to auto, and the flash was set to low power mode. Normally, the Bushnell™ immediately cuts power to the SD card once it has finished writing an image or images. This does not allow sufficient time for images to be transmitted from the WiFi SD card to the smart bridge using the WiFi network. To address this, the custom microcontroller keeps the WiFI SD card powered on until the images have been transmitted to the smart bridge.

The WiFi SD card is secured permanently into the camera (to prevent the power connection being damaged, Figure 3), but the images are also stored on a removable micro SD for later download if required.

**Figure 3.**
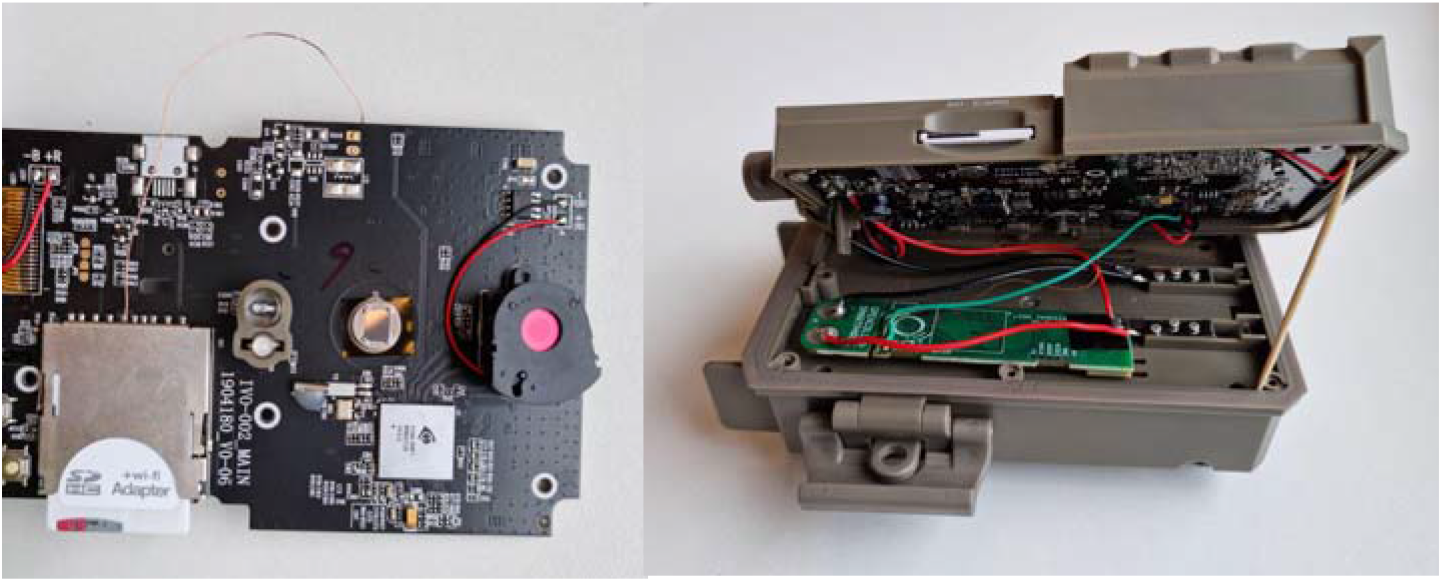
Modified bushnell™ camera trap showing the LoRA relay and printed circuit board, the WiFi SD card and power supply (removed and installed).

### Smart-bridge

The Smart-bridge (Figure 4) contains a custom printed circuit board (PCB) with a LoRa STM32L0 ultra-low-power microcontroller, RockBLOCK satellite modem and connections for a Raspberry Pi 4 Compute module. The hardware is stored in a weatherproof NANUK NANO 330 case (L188 × W130 × H65 mm). By default, the Raspberry Pi is turned off and thus the system consumes minimal power (less than 50 microampère; see *Power* later). When the smart bridge receives a LoRa message from a nearby camera it turns on the Raspberry Pi, which then downloads and classifies the images from the Bushnell™ using artificial intelligence. After sending the results over the satellite network (see later), the system powers down.

**Figure 4.**
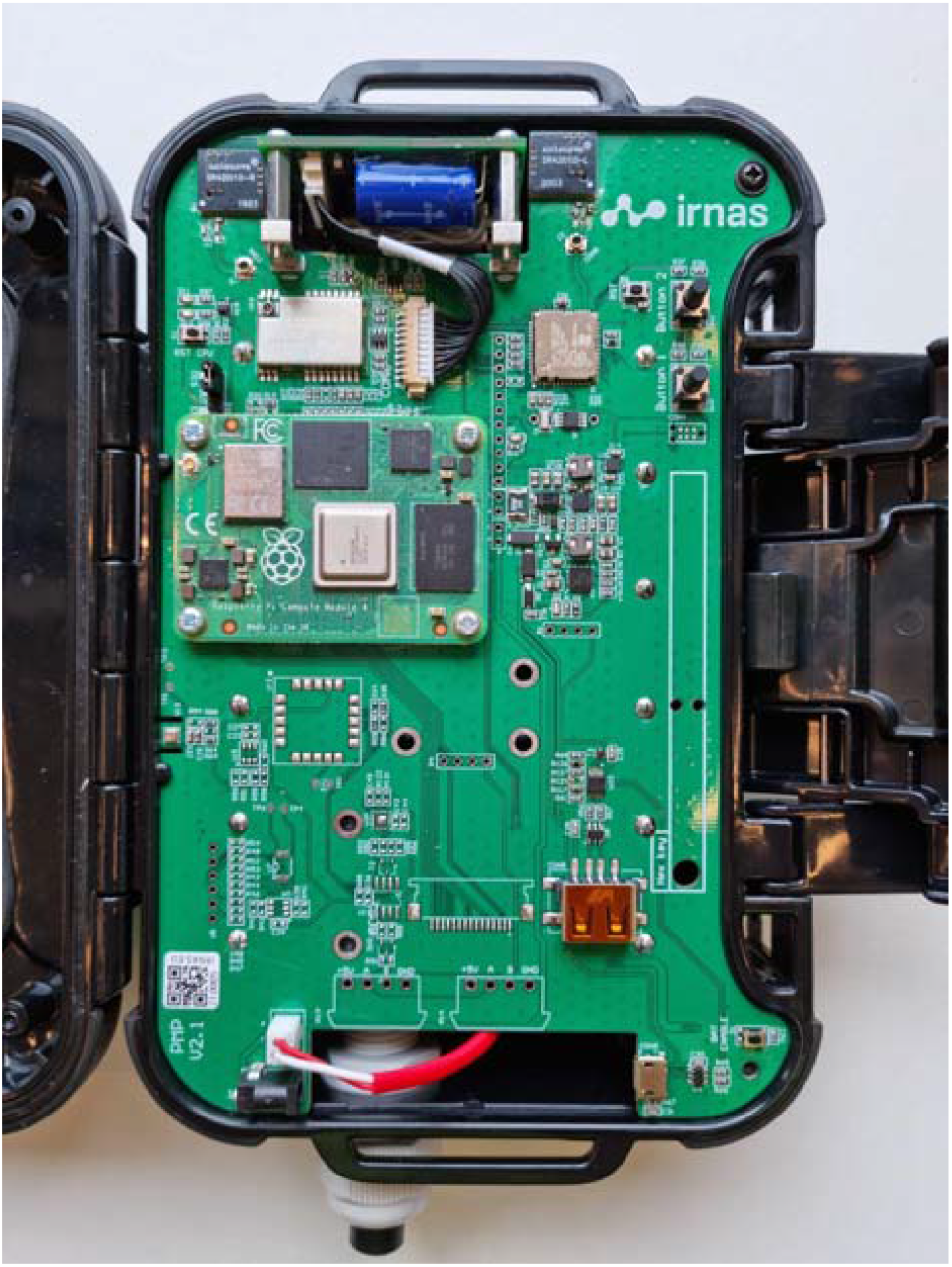
Smart Bridge PCB

### Raspberry Pi

The Raspberry Pi 4 compute module is integrated onto the Smart-bridge PCB with a pair of 100-pin mezzanine connectors. Raspberry Pis provide an excellent platform for development purposes and have been used widely in ecology (Jolles, 2021; Sethi et al., 2018; Sturley & Matalonga, 2020). Furthermore, although the Raspberry Pi 4 is power inefficient relative to other similar boards on the market (e.g. Arduino based systems), the Pi 4 can run artificial intelligence models built on relatively large architectures. Our approach of only briefly powering the Pi when needed allowed us to harness its computational power in an energy-optimised way.

The Raspberry Pi 4 Compute module runs Raspbian lite and Python 3 scripts together with the Tensorflow Lite runtime to fetch the images and run the artificial intelligence model. A SQLLite database is used to track image status (download status, transmission status etc).

### Satellite modem

There are many satellite networks available for civilian use. We chose the Iridium satellite network because it has near global coverage, is relatively inexpensive, and has widely available hardware including miniaturised, low-power modems. The Iridium network is also well known in the ecology community where it is regularly used for animal tracking using GPS collars. We used the RockBLOCK 9603 modem from Rock Seven to connect to the Iridium network.

### Environmental data

The smart bridge PCB is equipped with a temperature, humidity and barometric pressure sensor. Since these are mounted directly on the PCB they are not currently suitable for external environmental monitoring (other than barometric pressure) but they are useful for evaluating if the smart bridge is intact. For example, the smart bridge housing is completely sealed once closed and contains silica gel. In a humid environment such as a tropical forest, the humidity should drop once the bridge is installed and closed. A future rise in humidity could be used as an indicator of a possible hardware problem. We do not present data from these sensors or discuss them further.

### Power

For the smart bridge we used six NCR18650PF rechargeable batteries totalling 16,500 mAh power and a 6 volt 6 watt solar panel for charging. Initial testing in the Netherlands showed an active smart bridge, processing and transmitting approximately 17 images per day (see results), could be powered indefinitely by a solar panel without intervention.

For the Bushnell™ camera trap, we used six Energizer© Ultimate Lithium™ AA batteries (non-rechargeable). Normally the Bushnell™ has a battery life of approximately one year using these batteries. The addition of the microcontroller and the WiFi SD card draws additional power, however, which will reduce deployment times. During testing in the Netherlands the camera achieved three months of battery life when activated up to 17 times per day on average (see Results). We expect field deployment times to be longer than this since the camera is likely to be triggered less frequently when correctly installed and parameterized.

### Optimising alerts and minimizing data transmission costs

The Iridium satellite network supports short burst data and a maximum of 340 bytes can be sent in a single transmission. Satellite data is relatively expensive so we optimised the alerts to maximise the amount of information transmitted per message. The timestamp was reduced to 4 bytes by sending the number of elapsed seconds since January 1st 2010. The image label from the artificial intelligence model (e.g. elephant) was mapped to a 1 byte number and later converted back to a text label on the web backend. All other data, like AI prediction ‘confidence’ for the top-scoring species label (softmax algorithm probabilities), temperature and smart bridge voltage are mapped to 1 byte numbers. This allowed us to send up to 55 image classification results in a single satellite message.

## Software stack

### Artificial intelligence model

Our aim was to provide reliable alerts of species detections without requiring images to be transmitted to the end-user over a wireless network. Since our focus was on forest elephants during the pilot, we initially tested the model from (Whytock et al., 2021), which classifies 26 central African forest mammal and bird species, including forest elephants. However, the model was built using a relatively large convolutional neural network (CNN) architecture (ResNet50) and is 100 MB in size. This model took over 20 seconds to classify a single image using the Raspberry Pi 4 compute module, which drew a substantial amount of power and made the model unsuitable for our purposes.

To find a suitable alternative architecture to ResNet50, we compared inference times among a suite of 16 pre-trained computer vision models using their Fast.ai (Howard & Gugger, 2020) implementations (see Figure S1 for results). We did not evaluate classification accuracy using these models but only inference times. Then, we trained a Tensorflow Lite model (using Google Cloud’s AutoML service) and a Fast.ai model (SqueezeNet 1.1, the second-fastest from our tests) using a dataset of 105,000 images (a subset from Whytock et al. (2021)) with three, almost equally distributed classes (elephant, human and other). For these two models, we compared model precision and accuracy using a smaller, held out subset of 14,642 images, with almost equal distribution among the classes. We found that the TensorFlow Lite model provided the shortest inference time (∼100 ms vs ∼1200 ms for SqueezeNet) and precision and accuracy was similar between the two architectures (Table S1). Therefore, the Tensorflow Lite model trained using AutoML was chosen for deployment during the pilot.

### Back-end

An important element of receiving real-time alerts from camera traps is a centralised platform that can be used to receive, interpret and display the incoming data. Following our philosophy of using existing technology, we integrated the system with the EarthRanger platform (www.earthranger.org). Incoming data is first stored on our own Django-based back end. Once an alert is received the raw data is stored in a SQL database. A task-queue based system is then used to send the data to integrated platforms (e.g. EarthRanger or others). As well as offering a web-platform and mapping capabilities for displaying alerts, EarthRanger can also be configured to send messages in real-time using WhatsApp™, short message service (SMS), e-mail and other methods.

## Case study

Real-time alerts from cameras have many potential applications but our interest was testing if they could be used to help manage human-elephant interactions during crop depredation, in Gabon, central Africa. Gabon is almost 270,000 km^2^ with 88% of the country covered in closed-canopy forest. The country is home to more than 50% of the global population of the critically endangered forest elephant (Gobush et al., 2021). Although Gabon’s human population is relatively small (*c*. 2 million), with most people living in urban areas, rural communities across the country can suffer significant agricultural losses due to elephants (Walker, 2012). This affects the safety and wellbeing of both humans and elephants (e.g. retaliatory killing of elephants, humans injured or killed during interactions) and can have substantial economic consequences for rural communities (Terada, 2021).

Many villages work with Gabon’s National Park Agence (ANPN: the Agence Nationale des Parcs Nationaux) to manage elephant crop depredation. We therefore partnered with ANPN to test the camera’s ability to detect elephants and send real-time alerts to ANPN ecoguards (employees of the national park who lead fieldwork, tourism, and law enforcement) over WhatsApp™ in two locations. The first location was the Station d’Etudes des Gorilles et Chimpanzés (SEGC) in Lopé National Park, where elephants are common in the surrounding area. The facilities at the research station allowed us to test the system under controlled but realistic conditions (elephants regularly enter the station grounds). The second location was Kazamabika village, in the northern edge of Lopé National Park, where communities have established farms. Kazamabika received an electric fence to protect crops from elephants in 2016, and the local community is highly engaged in research to help understand and mitigate human-elephant conflict (Rakotonarivo et al., 2021). Although the electric fence is functional and effective, elephants still enter the village and surrounding forest to feed on domestic fruit trees that are also harvested by people. Although rare, elephants also occasionally succeed in entering the fence, potentially causing some damage to crops.

We tested whether alerts from the smart cameras could be used by ANPN ecoguards in Lopé National Park to detect when elephants are approaching the electric fence or village, allowing them to alert villagers to potential problems. There remains uncertainty about the most effective action villagers can take when they receive an alert, but at minimum they can have pre-warning and avoid the forest where elephants are detected to not be endangered, or they can take action to scare the elephants (e.g. creating noise, or smoke fires). In future, the system could potentially trigger auto-deterrents, such as sounds or lights, assuming effective deterrents are developed (see Discussion). Mitigating human-elephant conflict using sound, smoke, bees and plant species (e.g. chilli pepper) is an active area of research across Africa and Asia (Dror et al., 2020; Ndlovu et al., 2016; Pozo et al., 2019) and we did not explore the effectiveness of particular deterrents during our trials.

### Field testing

We constructed seven systems and tested five under different settings for a combined total of 72 days (Table 1). Camera locations were chosen to test (a) how the position of the smart bridge and vegetation structure (e.g. forest canopy cover) affected data transmission and satellite connectivity, (b) how far the smart bridge could be installed from the camera, (c) how well the solar panel functioned under different light levels, and (d) how well the artificial intelligence algorithm performed with different camera backgrounds (open areas, farmland and forest). We chose the testing locations based on qualitative differences in vegetation structure, light availability and image background (Table 1). In summary, the smart bridge and solar panel were installed together on a tree 2 - 6 m above ground level at a distance of 5 - 20 m from the camera trap. Camera traps were installed on a tree approximately 40 - 50 cm above ground level, perpendicular to and approximately four metres from the centre of well-used elephant paths.

**Table 1.**
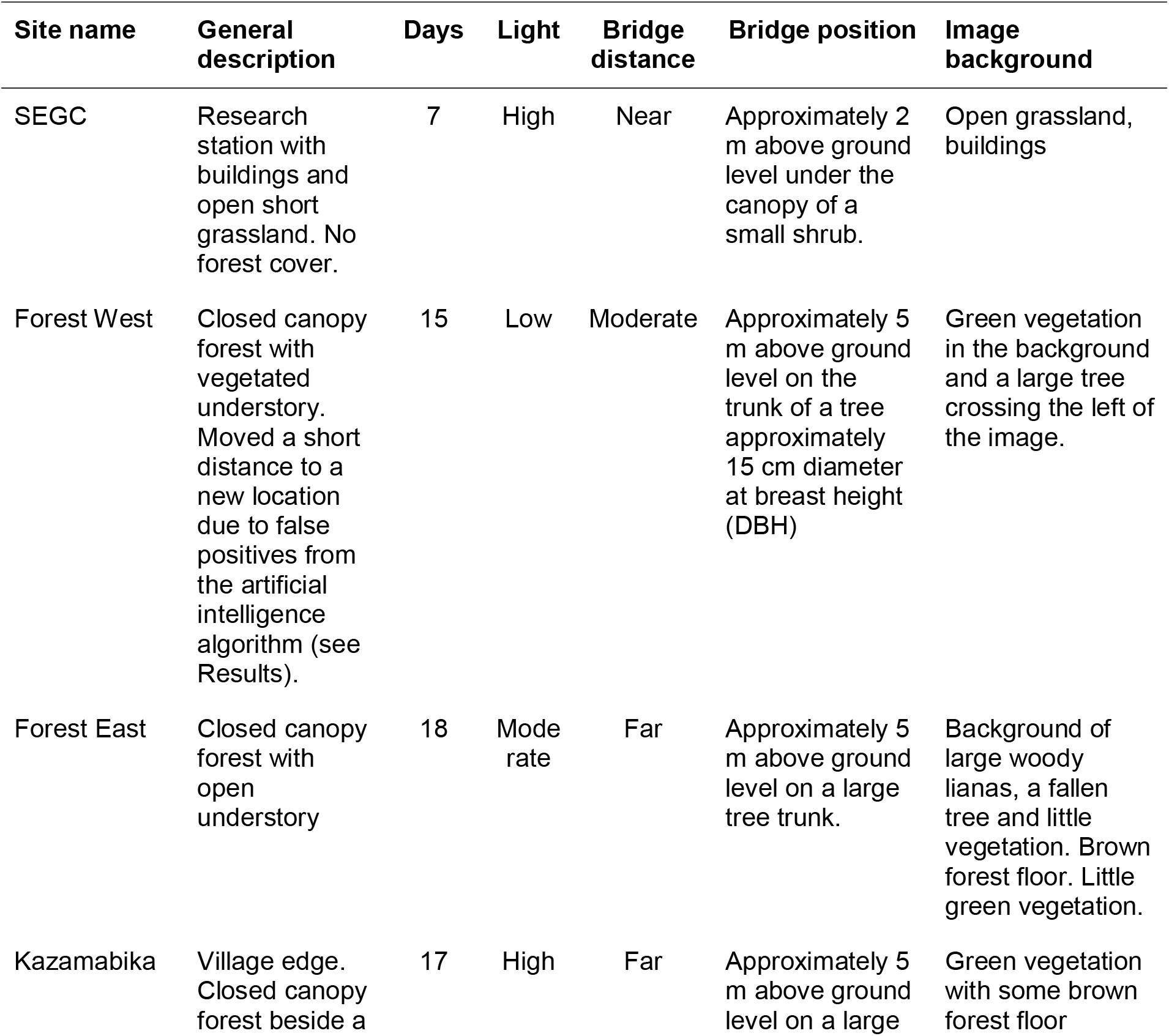
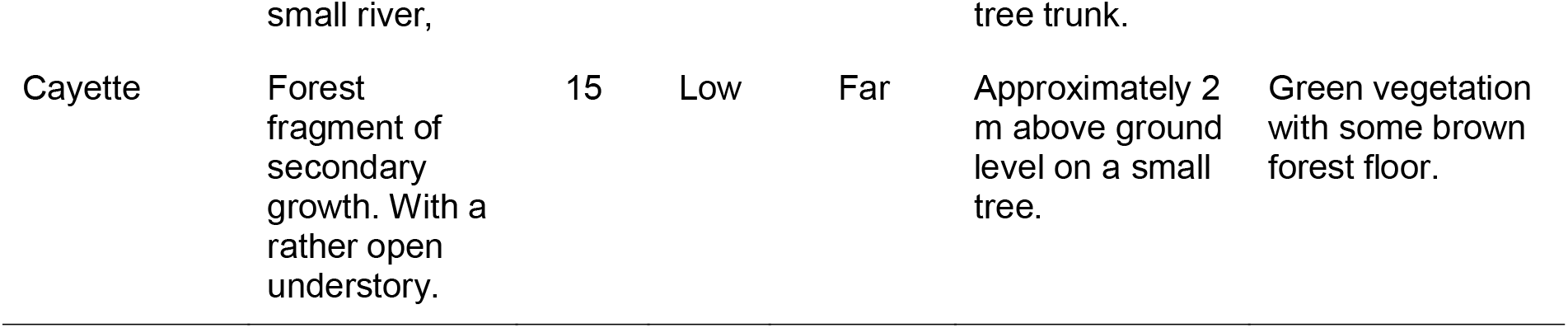
Description of test locations and field conditions with qualitative descriptions of light availability (Light: low, medium, high), distance between camera and smart bridge (Bridge: near < 5 m, moderate 5 - 10 m, far 10 - 20 m), the positioning of the Smart Bridge (Bridge position) and image background (considered important for artificial intelligence performance).

We compared results from field testing with benchmark data from two systems operated in the Netherlands for three months during the development stage. Both of these systems were deployed in urban settings (a private garden and empty roof-top) with a clear view of the sky. During field testing, all images were stored on the camera trap SD card and retrieved at the end of the testing period for validating artificial intelligence labels.

### Data analysis

To evaluate the speed at which alerts were transmitted and received, we calculated the median time-difference in minutes between image capture and receipt of the alert by the back end for each location individually, and for all stations. For each of the test locations we also created time-series plots showing changes in smart bridge power during deployment. Camera power was also monitored during tests in the Netherlands but not during the field testing.

We assessed artificial intelligence model performance (precision, recall, accuracy and F1 score (Kuhn, 2020)) on the newly captured images by comparing artificial intelligence-generated image labels with ‘expert’ labels. Expert labels were created by first labeling the captured images using the Mbaza AI software (Whytock et al., 2021) and manually validating all results (co-author RW).

During field testing we observed that, within a given image sequence of elephants (i.e. a number of images taken during the same presence event), the first and last images could be mislabelled when only a small part of the elephant was visible. We therefore tested if (a) a simple vote-counting approach (i.e. counting the most frequently predicted top-one label in an image series) could improve predictions on an event, and (b) if thresholding on the softmax values (i.e. excluding images below a softmax threshold before vote counting) could improve event prediction accuracy. Events were defined as a series of images taken within an independent 30-minute time window. Softmax thresholds were from 0 to 0.9 in 0.1 intervals. In some instances, vote counting resulted in a tie between the number of votes for each class. In these cases, we chose ‘elephant’ if it was among the ties, or otherwise chose the label ‘other’.

## Results

A total of 814 images were captured during the field test (Table S2) and alerts for 588 images were received by the backend. Of the 226 alerts not received, 72 were from Cayette, which was not able to send any alerts due to the position of the smart bridge (2 m above ground level under a tall, closed canopy) and 154 were from Forest East because the smart bridge unexpectedly ran out of battery after just six days. This was caused by a problem with the charging circuit and was inconsistent with tests in the Netherlands, which achieved > 3 months of battery life (see *Battery life* for further details). We removed a further 17 images which had no timestamp (human error during camera setup) and which could not be used to evaluate alert time delays, leaving *n* = 571 alerts from four systems for the analysis.

### Alert times

There was a median 7.35 minutes time difference between capturing an image and sending an alert (*n* = 4 camera stations). Median, minimum and maximum alert times are given in Table S3 for each location. Of the four systems, Kazamabika had the slowest median alert time (306.3 min). A total of 296 (52%) of alerts were received within 15 minutes or less (Figure 5, Figure S2).

**Figure 5.**
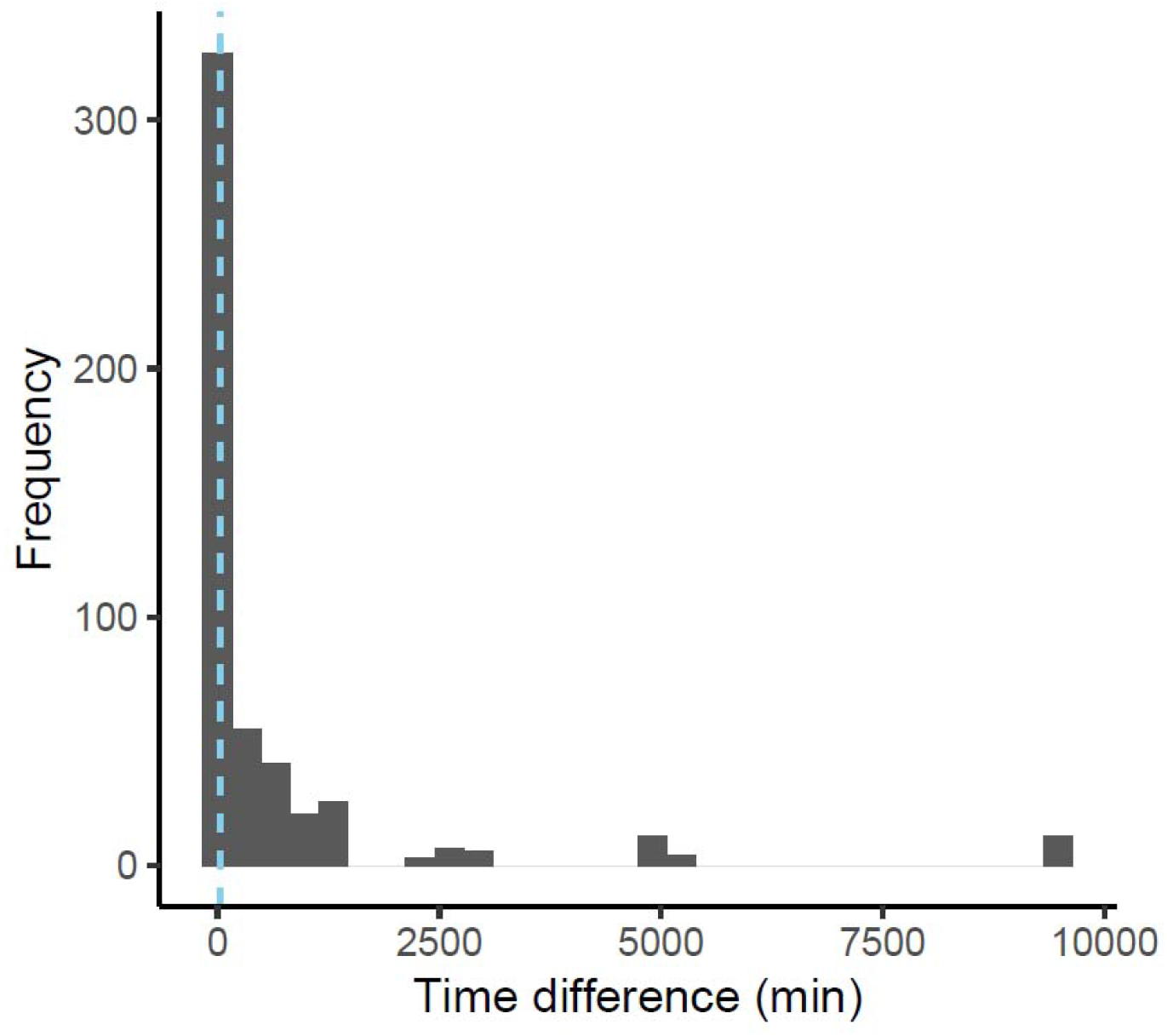
Histogram showing time difference between image capture and alert transmission time. The dashed line shows the median alert time of 7.35 minutes.

### Battery life

Preliminary tests in the Netherlands showed that even with a median of 17.23 image captures per day (range 0 - 40), the systems could operate continuously during the winter under low sunlight for a minimum of three months (Figure 6). During field testing in Gabon, we found mixed results (Figure 7) and one system discharged in six days (Forest East). Forest West lasted the full 18 days but did not show signs of substantial charging as was seen in the Netherlands. Kazamabika and SEGC both operated as expected.

**Figure 6.**
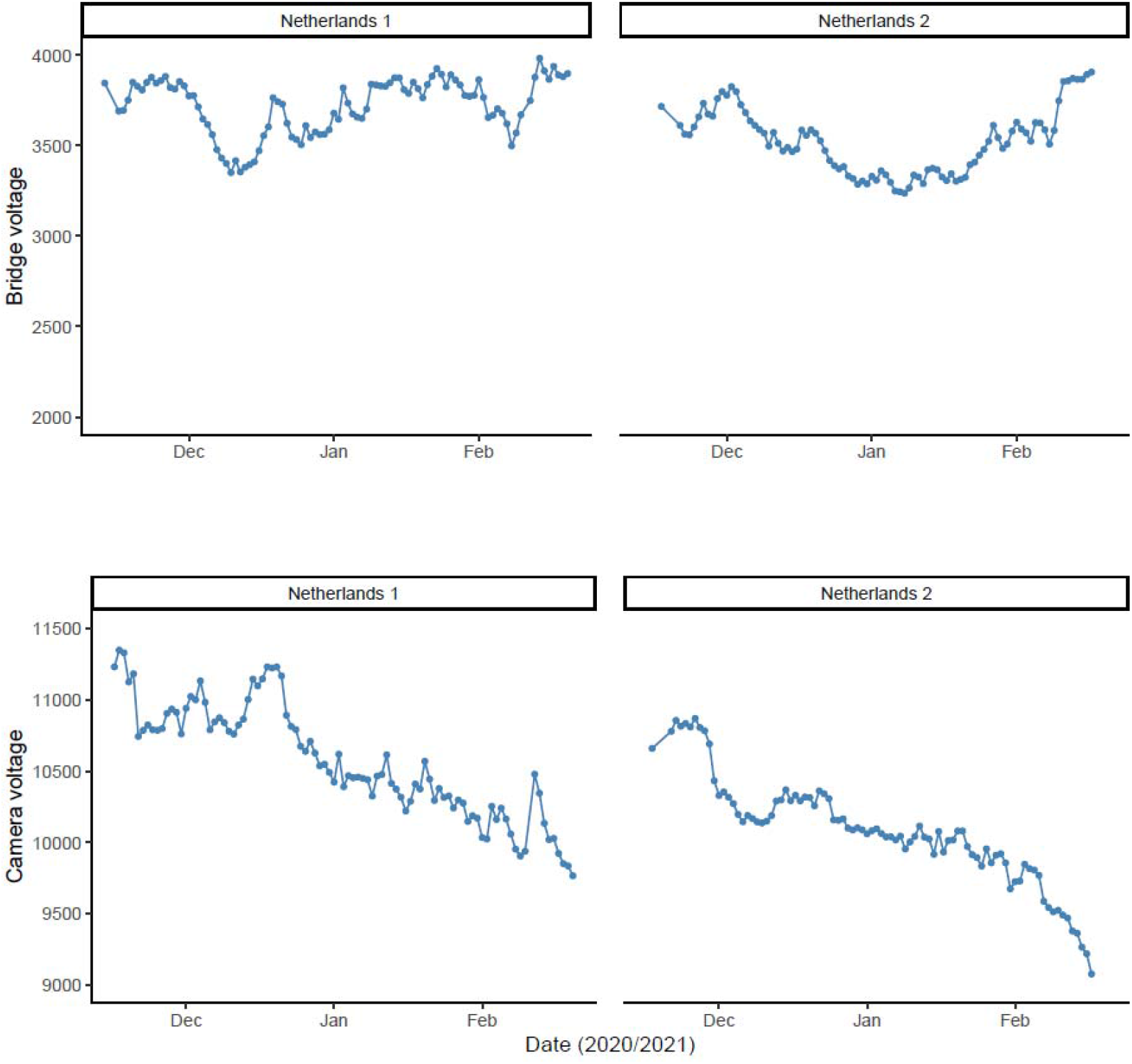
Smart bridge and camera voltage change over time during testing of two systems in the Netherlands using a solar panel.

**Figure 7.**
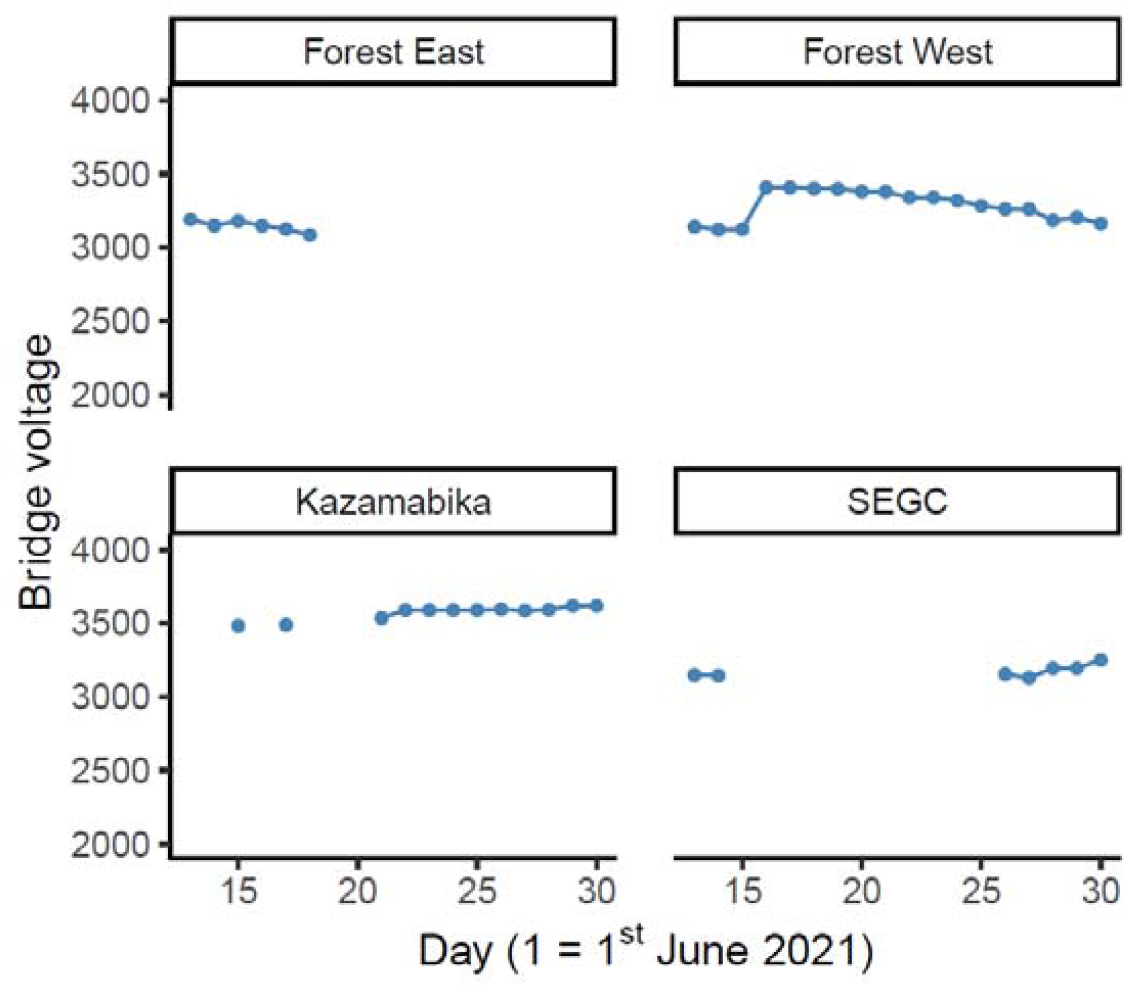
Smart bridge voltage changes over time during testing of four systems in Gabon using a solar panel.

Initially it was thought that the forest canopy was preventing charging by the solar panel in Forest East and Forest West, despite careful positioning. However, further tests revealed the mechanism designed to prevent the charging circuit from overheating was being triggered prematurely by the high ambient temperatures and high voltage output from the solar panel in Gabon, in contrast to the Netherlands. This problem has been solved by removing the overheating protection.

### Artificial intelligence model accuracy and interpreting alerts

Overall model accuracy on new data collected during the field test (*n* = 571 images) was 84%, with a Kappa statistic of 0.74. For the elephant class, precision was 82% and recall 86%, with a balanced accuracy of 86%. Test statistics for all classes and a confusion matrix are given in Table 2 and Figure 8. Classification of events using vote counting without any softmax thresholding (i.e. choosing the most frequently predicted class in a 30 minute time window) gave an overall performance of 78% and a Kappa statistic of 0.64 (*n* = 142 events) (Table 2). Excluding uncertain image labels using a softmax threshold before vote counting improved overall accuracy for event classification, as well as balanced accuracy for the elephant events (*n* = 29 true events, *n* = 30 predicted), which reached 98% at a threshold where images were excluded with a softmax value < 0.9 (Figure 9). This almost matched human accuracy with just one false positive event and no false negatives.

**Table 2.** Model performance by class for *n* = 571 images and *n* = 142 events using vote

| Test statistic | Elephant_African | Human | Other |
| --- | --- | --- | --- |
| *Images* |  |  |  |
| Pos Pred Value | 0.82 | 0.92 | 0.81 |
| Neg Pred Value | 0.89 | 0.95 | 0.91 |
| Precision | 0.82 | 0.92 | 0.81 |
| Recall | 0.86 | 0.80 | 0.82 |
| F1 | 0.84 | 0.86 | 0.82 |
| Prevalence | 0.44 | 0.22 | 0.34 |
| Detection Rate | 0.38 | 0.18 | 0.28 |
| Detection Prevalence | 0.46 | 0.20 | 0.34 |
| Balanced Accuracy | 0.86 | 0.89 | 0.86 |
| *Events* |  |  |  |
| Pos Pred Value | 0.62 | 0.72 | 0.94 |
| Neg Pred Value | 0.97 | 0.96 | 0.72 |
| Precision | 0.62 | 0.72 | 0.94 |
| Recall | 0.90 | 0.81 | 0.77 |
| F1 | 0.73 | 0.76 | 0.85 |
| Prevalence | 0.20 | 0.18 | 0.61 |
| Detection Rate | 0.18 | 0.15 | 0.47 |
| Detection Prevalence | 0.30 | 0.20 | 0.50 |
| Balanced Accuracy | 0.88 | 0.87 | 0.85 |

**Figure 8.**
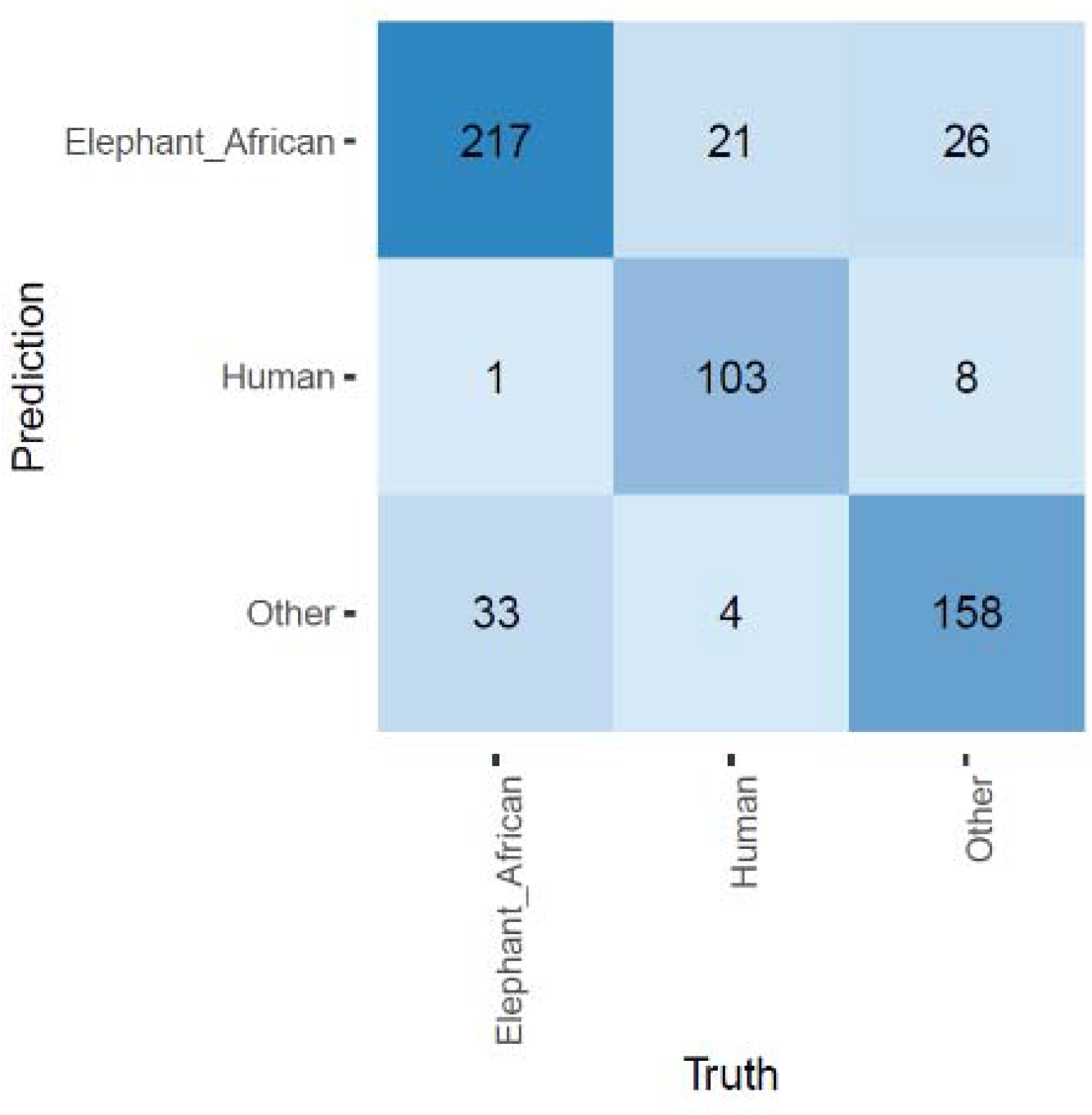
Confusion matrix for image-based classification

**Figure 9.**
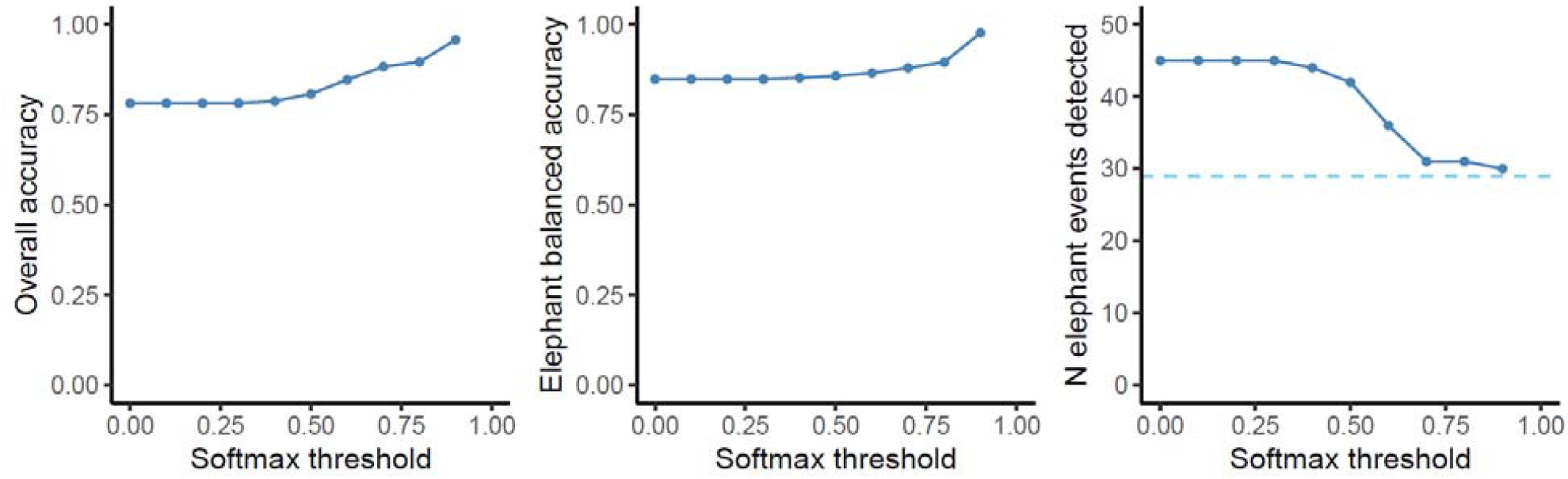
Effects of using a softmax threshold to exclude uncertain labels before vote counting to classify an event on (a) overall accuracy, (b) balanced accuracy for events labelled as elephant and (c) the number of elephant events detected (dashed horizontal line shows *n* = 29 true events).

One camera (Forest West) returned several false-positive elephant detections during the first two days of deployment. Verification of the images in the field showed this was likely to be caused by an unusual branch resembling an elephant trunk or limb, close to the camera lens. Moving the camera to another location a short distance away solved this issue.

## Discussion

Sending real-time alerts from ecological sensors such as camera traps in areas with poor data connectivity is complex and involves fine tuning a large number of software and hardware parameters. These include camera settings, camera positioning, achieving reliable network connectivity, training and running artificial intelligence models, interpreting and displaying artificial intelligence outputs and providing a reliable source of power. Our results demonstrate that these parameters can be tuned to achieve reliable, near real-time alerts from camera traps under challenging field conditions. We also identified potential pitfalls and areas that should be prioritised for future research and development.

### Problems and solutions

Battery charging using the solar panel in Gabon did not function in forests as expected given results from testing in the Netherlands. However, this was rapidly diagnosed as an issue with the charging circuit and has now been rectified.

At one camera location, false positive elephant detections were quickly remedied by moving the camera position. However, this could be difficult to detect during a real deployment after cameras have been left in-situ by field teams. Improved models and training data will likely reduce this issue in future (Beery et al., 2018). The problem can also be mitigated by ensuring that cameras are positioned so that new images replicate training data as closely as possible.

A total of 588 alerts were generated by our four systems during 18 days of testing, and the final total could have been as high as 814 if all alerts had been received. This is a substantial amount of data to interpret on a rolling basis with just four systems and three label classes. In future, we recommend first implementing vote-counting combined with softmax thresholding on the smart bridge to reduce the total number of alerts, which would have been just 30 (with one false positive) if restricted only to elephants. Similar vote counting approaches have also been successfully used to summarise camera trap image labels made by citizen scientists using online platforms (Swanson et al., 2015).

Summarising alerts into temporally independent events using vote counting would not only improve alert accuracy but also reduce data transmission costs. This approach will be implemented into future versions of the smart camera system.

Our system does not currently send images but this would be possible using an on-demand approach. For example, users could request certain images or an image series by sending a message (relayed via satellite) to the smart bridge. The main limitations to implementing this is achieving a reasonable trade-off between image quality and transmission cost. For example, sending an extremely compressed thumbnail would cost $2 USD per image with a $20 per month contract on the Iridium network (Figure 10). Scaling this up to hundreds of cameras could be financially unfeasible for many use-cases.

**Figure 10.**
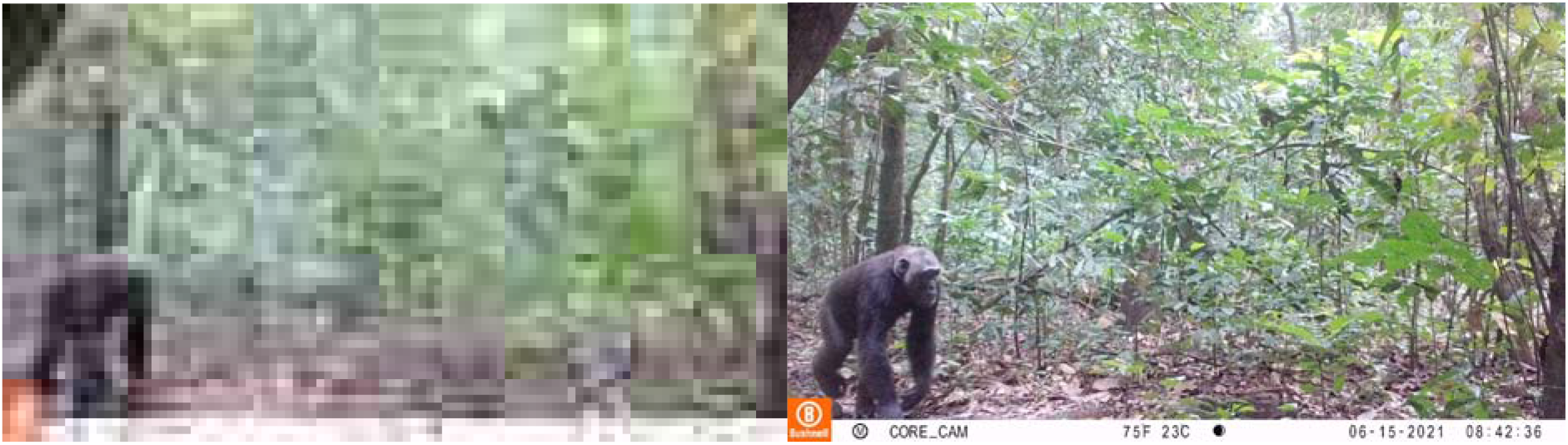
Camera trap image of a chimpanzee, with example compressed thumbnail (left) compared to the original image (right). The compressed thumbnail would require three messages sent over the Iridium network using a RockBlock modem and cost approximately 2 USD (on a 20 USD monthly contract). The thumbnail provides limited information for interpretation by both human and artificial intelligence algorithms.

The next generation of camera traps will run artificial intelligence models on the camera hardware directly (known as ‘edge computing’) instead of using a separate smart bridge. However, if the goal is to transmit real-time data from cameras installed near the ground for wildlife monitoring, then developers should be aware that it will be difficult to achieve network connectivity under a dense forest canopy. We were not able to send any images from Cayette forest patch, where the smart bridge was installed just 2 m above ground level. The wireless smart bridge, which can be mounted in a tree, might therefore be a useful design feature for future edge computing solutions.

A final problem that only became apparent during field testing was that users need to know if the system is still functioning when no alerts are received. The latest version of the system now sends a timed, daily ‘keep-alive’ message notifying the user that it is functioning as expected.

### Potential applications beyond our case study

Our results show that we have created a viable hardware solution for running powerful artificial intelligence algorithms in the field and transmitting results over a satellite network. The computing power of the Raspberry Pi 4 is currently underused and there is scope for attaching other sensors, such as microphones for bioacoustic recording.

There are already a substantial number of open-source Raspberry Pi projects available for ecological research, and many of these could be integrated with the smart bridge with relatively minimal effort (Jolles, 2021). Likewise, there is scope for implementing other artificial intelligence models, for example to count animals in images or to recognise species from other ecoregions. The list of potential applications for the hardware is limited only by imagination, but some examples relevant to ecology and conservation are given in Table 3.

**Table 3.**
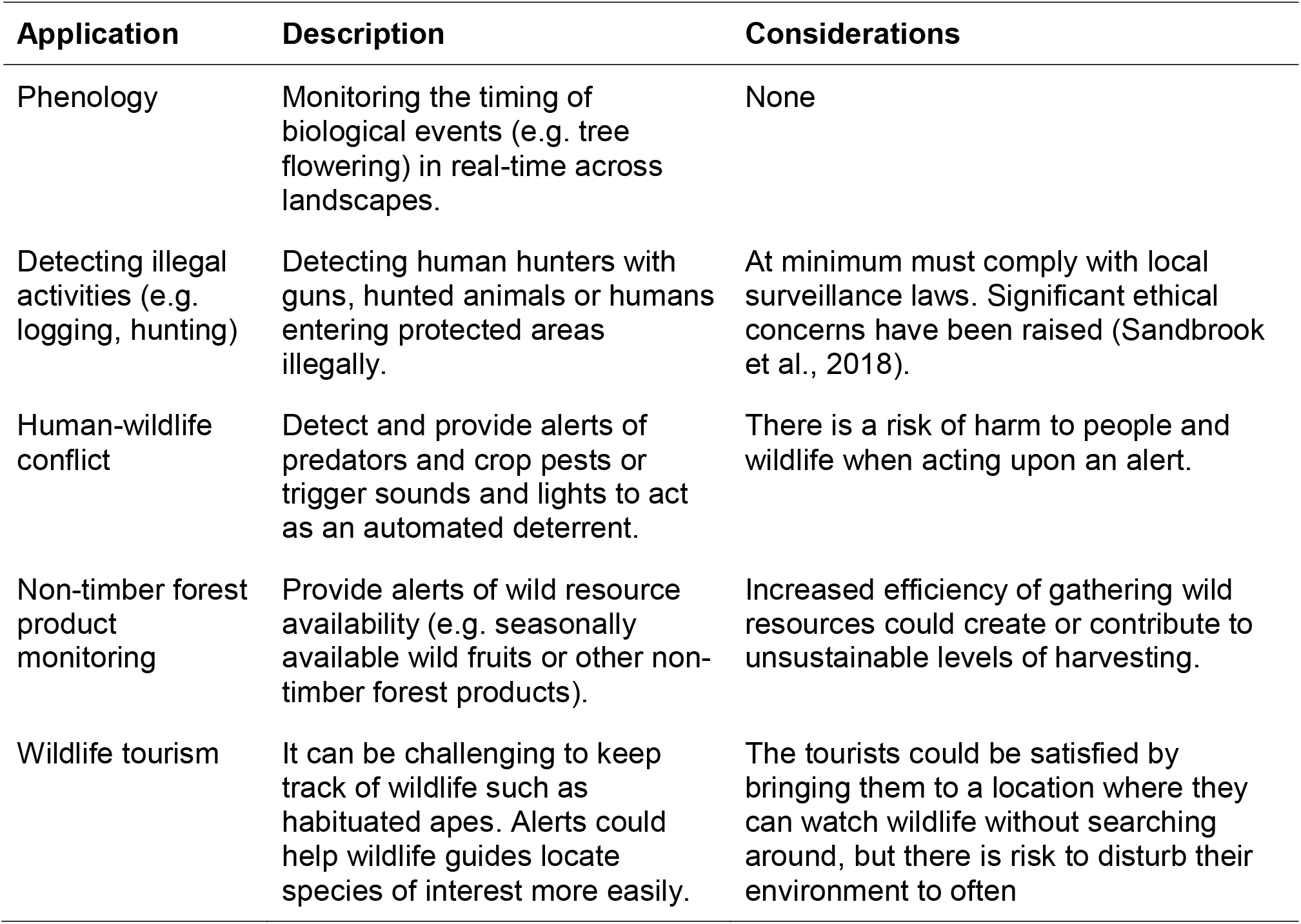
Potential ecology and conservation applications for real-time, artificial intelligence-enabled smart cameras

| Application | Description | Considerations |
| --- | --- | --- |
| Phenology | Monitoring the timing of biological events (e.g. tree flowering) in real-time across landscapes. | None |
| Detecting illegal activities (e.g. logging, hunting) | Detecting human hunters with guns, hunted animals or humans entering protected areas illegally. | At minimum must comply with local surveillance laws. Significant ethical concerns have been raised (Sandbrook et al., 2018). |
| Human-wildlife conflict | Detect and provide alerts of predators and crop pests or trigger sounds and lights to act as an automated deterrent. | There is a risk of harm to people and wildlife when acting upon an alert. |
| Non-timber forest product monitoring | Provide alerts of wild resource availability (e.g. seasonally available wild fruits or other non-timber forest products). | Increased efficiency of gathering wild resources could create or contribute to unsustainable levels of harvesting. |
| Wildlife tourism | It can be challenging to keep track of wildlife such as habituated apes. Alerts could help wildlife guides locate species of interest more easily. | The tourists could be satisfied by bringing them to a location where they can watch wildlife without searching around, but there is risk to disturb their environment to often |

## Application Description Considerations

### Current limitations

Using the system outside of our case study would require both technical expertise to build or modify all of the necessary hardware components and sufficient training data to create a new artificial intelligence model. The Audiomoth bioacoustic recorder (Hill et al., 2018) project has overcome this challenge using a ‘group buy’ format, where the design is completely open-source and customers order the units in advance. The units are then only manufactured and shipped when a target number is reached. Currently, the system presented here costs approximately 1000 euros per unit including the camera, smart bridge and solar panel, but this does not include labour costs for building the units, satellite contract costs or field deployments. This is more expensive than a standard camera trap but like all technology these costs will reduce in future. We anticipate that our approach will be superseded by new developments in the next five years, but hope the lessons learned here can help drive and inform the development of new technologies.

Other limitations include the sometimes low accuracy of the artificial intelligence model at the image-level. However, our main focus was building a complete system that was field-ready rather than attempting to achieve perfect artificial intelligence predictions, and we found that the model was usable, particularly when applying a vote-counting approach. Improved models can be built using new incoming data and new approaches will give gains in precision and accuracy in future (Beery et al., 2019; Schneider et al., 2019).

## Conclusion

We have shown that it is possible to send reliable, real-time information from camera traps over the Iridium satellite network by integrating artificial intelligence, off-the-shelf and custom hardware. Our solution does not depend on installation of additional network infrastructure in the landscape and can be operated by non-experts from anywhere on earth. Real-time data gathering and interpretation will change how ecologists and conservationists understand and manage ecosystems. We piloted the system for detecting elephants, but new artificial intelligence algorithms will be created in future to capture other species or objects in images, such as illegal human activities in protected areas.

## Acknowledgements

Q42 funded hardware research and development costs. RCW was funded by the EU 11th FED ECOFAC6 program grant to the National Parks Agency of Gabon during data collection and curating of images used to train the artificial intelligence model, RCW, DMI and KAA received funding from the UK Research and Innovation’s Global Challenges Research Fund (UKRI GCRF) through the Trade, Development and the Environment Hub project (project number ES/S008160/1) during the field trials reported here. DL, NM, LM and BM were funded by the EU 11th FED ECOFAC6 program grant to the National Parks Agency of Gabon throughout all aspects of the study. The Ministry of the Environment, Water and Forests funded LJTW and co-funded fieldwork in Gabon. Appsilon Data Science funded the artificial intelligence model development costs. We thank Kelly Boekee and Cisquet Kiebou Opepa for camera trap data made available by the Tropical Ecology Assessment and Monitoring Network (now https://wildlifeinsights.org). AC was funded by the Hertford College Mortimer May Fund at Oxford University. KAA and RCW received funding from forestLAB during the writing of this paper. We thank Smart Parks for the ability to build upon their open-source hardware designs and Irnas for the hardware development. We thank Hugh Robinson and Ross Pitman for insights during early discussions.

## Author contribution statement

RCW contributed to the system design, co-wrote the manuscript, collected the data and analysed the data. TS designed the system, co-wrote the manuscript and collected data. TvD co-designed the system and collected data. JS created the AI model. HM co-designed the pilot. NM collected data. JAZ supplied data for the AI model. AFKP supervised RCW and contributed to writing the manuscript. LB supplied data for the AI model. SB supplied data for the AI model and contributed to writing the manuscript.

AWC supplied data for the AI model. PH supplied data for the AI model and contributed to the system’s design. DL contributed to the system’s design and contributed to writing the manuscript. BM supplied data for the AI model and collected data. LM collected data. CO contributed to the system’s design and supplied data for the AI model. LJTW contributed to the system’s design and co-designed the pilot study. DMI contributed to manuscript writing and interpreting results. KAA contributed to the manuscript, contributed to the system’s design and co-designed the pilot.

## Ethics statement

The work was approved by the University of Stirling General University Ethics Panel, application number GUEP (2021) 1044.

## Research permissions

The work was carried out in collaboration with the Tropical Ecology Research Institute in Gabon as part of the GCRF-TRADE Hub partnership.

## Data availability statement

*All data used in the analyses (excluding raw images) will be made publicly available on acceptance of the manuscript*.

## Supplementary Material

**Table S1.** Comparison between model accuracy for the Fast.ai SqueezeNet model and the TensorFlow Lite model trained using three classes.

| Model | Class | Number correct | Percentage correct |
| --- | --- | --- | --- |
| SqueezeNet | Human | 4529 / 5000 | 90.58% |
| TensorFlow Lite | Human | 4631 / 5000 | 92.62% |
| SqueezeNet | Elephant_African | 4516 / 5000 | 90.32% |
| Tensorflow Lite | Elephant_African | 4507 / 5000 | 90.14% |
| SqueezeNet | Other | 4358 / 5000 | 87.16% |
| Tensorflow Lite | Other | 4586 / 5000 | 91.72% |

**Table S2.** Images captured during each day of the field test for each location. NA indicates the system was deactivated.

| Day (June 2021) | Forest East | Forest West | Cayette | Kazamabika | SEGC |
| --- | --- | --- | --- | --- | --- |
| 13 | 12 | 2 | NA | NA | 50 |
| 14 | 18 | 4 | NA | 24 | 5 |
| 15 | 13 | 14 | NA | 1 | NA |
| 16 | 13 | 29 | 5 | 5 | NA |
| 17 | 8 | 4 | 4 | 32 | NA |
| 18 | 17 | 5 | 0 | 0 | NA |
| 19 | 1 | 8 | 3 | 0 | NA |
| 20 | 10 | 12 | 1 | 0 | NA |
| 21 | 7 | 9 | 11 | 32 | NA |
| 22 | 1 | 8 | 7 | 13 | NA |
| 23 | 16 | 30 | 2 | 1 | NA |
| 24 | 7 | 2 | 7 | 2 | NA |
| 25 | 12 | 7 | 7 | 27 | NA |
| 26 | 8 | 40 | 2 | 2 | 8 |
| 27 | 38 | 18 | 7 | 18 | 4 |
| 28 | 12 | 34 | 2 | 1 | 4 |
| 29 | 25 | 1 | 3 | 6 | 4 |
| 30 | 4 | 27 | 13 | 3 | 5 |

**Table S3.** Mean, minimum and maximum time difference between image creation time and alert time for four sites.

| Site | Minutes | Min. | Max. |
| --- | --- | --- | --- |
| Forest East | 6.9 | 2.37 | 863.8 |
| Forest West | 6.5 | 1.28 | 1299.2 |
| Kazamabika | 306.3 | 1.68 | 9473.9 |
| SEGC | < 1 | < 1 | 1277.1 |

**Figure S1.**
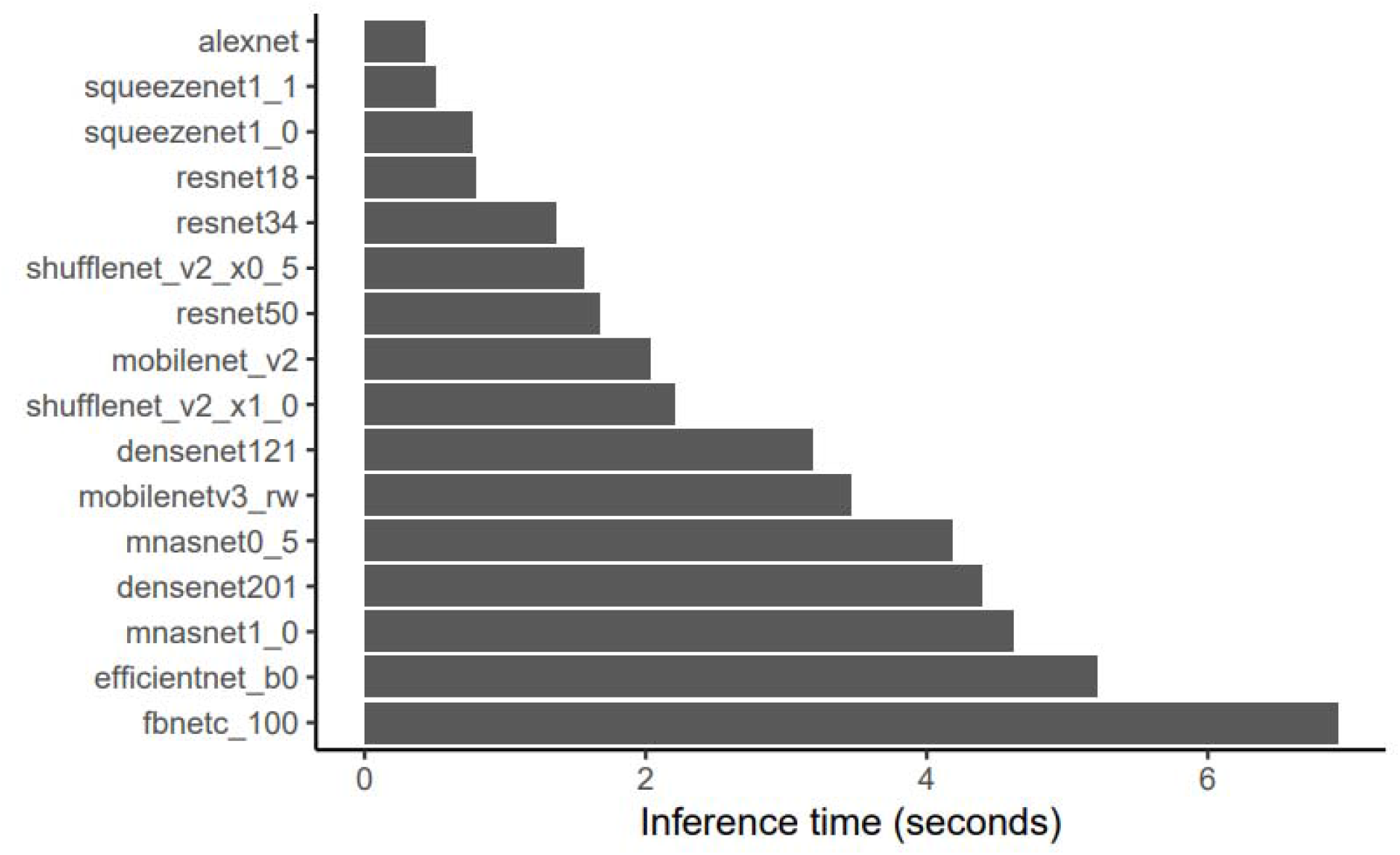
Mean inference time in seconds (*n* = 8 images 224 × 224 pixels) for 16 pre-trained computer vision CNN architectures run on the Raspberry Pi 4 compute module using PyTorch.

**Figure S2.**
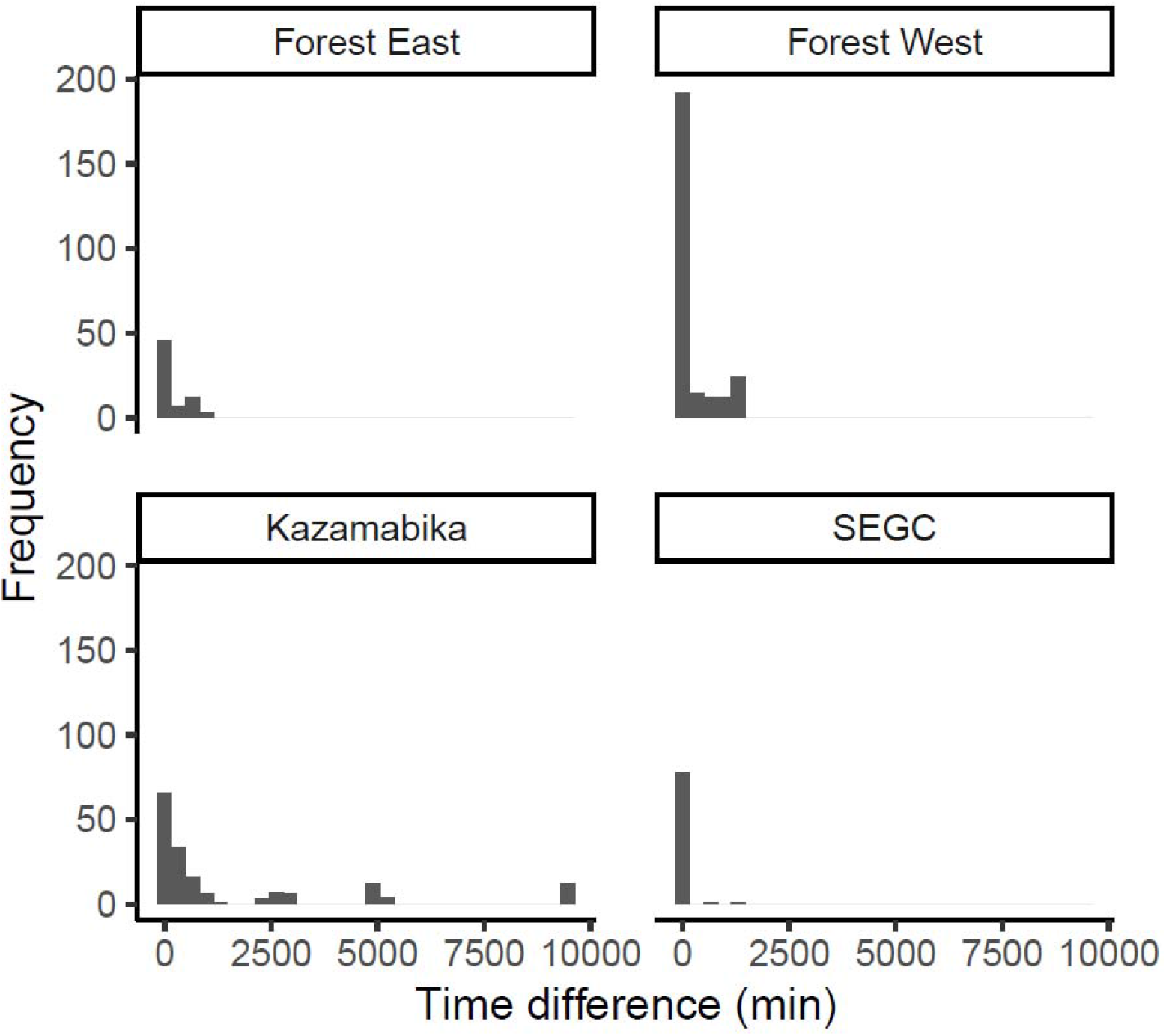
Alert times for each of *n* = 4 camera stations.

## References

Beery, S., Morris, D., Yang, S., Simon, M., Norouzzadeh, A., & Joshi, N. (2019). Efficient Pipeline for Automating Species ID in new Camera Trap Projects. Biodiversity Information Science and Standards, 3, e37222. https://doi.org/10.3897/biss.3.37222

Beery, S., Van Horn, G., & Perona, P. (2018). Recognition in Terra Incognita. Proceedings of the European Conference on Computer Vision (ECCV), 456–473. https://openaccess.thecvf.com/content_ECCV_2018/html/Beery_Recognition_in_Terra_ECCV_2018_paper.html

Bessone, M., Kühl, H. S., Hohmann, G., Herbinger, I., N’Goran, K. P., Asanzi, P., Costa, P. B. D., Dérozier, V., Fotsing, E. D. B., Beka, B. I., Iyomi, M. D., Iyatshi, I. B., Kafando, P., Kambere, M. A., Moundzoho, D. B., Wanzalire, M. L. K., & Fruth, B. (2020). Drawn out of the shadows: Surveying secretive forest species with camera trap distance sampling. Journal of Applied Ecology, 57(5), 963–974. https://doi.org/10.1111/1365-2664.13602

Convention on Biological Diversity. (2021). Report of the Open-ended Working Group on the Post-2020 Global Biodiversity Framework on its third meeting (Part I). 167.

Droissart, V., Azandi, L., Onguene, E. R., Savignac, M., Smith, T. B., & Deblauwe, V. (2021). PICT: A low-cost, modular, open-source camera trap system to study plant–insect interactions. Methods in Ecology and Evolution, 12(8), 1389–1396. https://doi.org/10.1111/2041-210X.13618

Dror, S., Harich, F., Duangphakdee, O., Savini, T., Pogány, Á., Roberts, J., Geheran, J., & Treydte, A. C. (2020). Are Asian elephants afraid of honeybees? Experimental studies in northern Thailand. Mammalian Biology, 100(4), 355–363. https://doi.org/10.1007/s42991-020-00042-w

Gobush, K. S., Edwards, C. T. T., Maisels, F., Wittemyer, G., Balfour, D., & Taylor, R. D. (2021). Loxodonta cyclotis. The IUCN Red List of Threatened Species 2021. https://dx.doi.org/10.2305/IUCN.UK.2021-1.RLTS.T181007989A204404464.en

Hill, A. P., Prince, P., Covarrubias, E. P., Doncaster, C. P., Snaddon, J. L., & Rogers, A. (2018). AudioMoth: Evaluation of a smart open acoustic device for monitoring biodiversity and the environment. Methods in Ecology and Evolution, 9(5), 1199–1211. https://doi.org/10.1111/2041-210X.12955

Hobbs, M. T., & Brehme, C. S. (2017). An improved camera trap for amphibians, reptiles, small mammals, and large invertebrates. https://doi.org/10.1371/journal.pone.0185026

Howard, J., & Gugger, S. (2020). Fastai: A Layered API for Deep Learning. Information, 11(2), 108. https://doi.org/10.3390/info11020108

Jolles, J. W. (2021). Broad-scale applications of the Raspberry Pi: A review and guide for biologists. Methods in Ecology and Evolution, n/a(n/a). https://doi.org/10.1111/2041-210X.13652

Kuhn, M. (2020). caret: Classification and Regression Training. R Package Version 6.0-86. https://CRAN.R-project.org/package=caret

Leidig, M., & Teeuw, R. M. (2015). Quantifying and Mapping Global Data Poverty. PLOS ONE, 10(11), e0142076. https://doi.org/10.1371/journal.pone.0142076

Ndlovu, M., Devereux, E., Chieffe, M., Asklof, K., & Russo, A. (2016). Responses of African elephants towards a bee threat: Its application in mitigating human-elephant conflict. South African Journal of Science, 112(1–2), 01–05. https://doi.org/10.17159/sajs.2016/20150058

Norouzzadeh, M. S., Nguyen, A., Kosmala, M., Swanson, A., Palmer, M. S., Packer, C., & Clune, J. (2018). Automatically identifying, counting, and describing wild animals in camera-trap images with deep learning. Proceedings of the National Academy of Sciences, 115(25), E5716–E5725. https://doi.org/10.1073/pnas.1719367115

Pozo, R. A., Coulson, T., McCulloch, G., Stronza, A., & Songhurst, A. (2019). Chilli-briquettes modify the temporal behaviour of elephants, but not their numbers. Oryx, 53(1), 100–108. https://doi.org/10.1017/S0030605317001235

Rakotonarivo, S., Bell, A., Abernethy, K., Minderman, J., Duthie, A., Redpath, S., Keane, A., Travers, H., Bourgeois, S., Moukagni, L.-L., Cusack, J., Jones, I., Pozo, R., & Bunnefeld, N. (2021). The role of incentive-based instruments and social equity in conservation conflict interventions. Ecology and Society, 26(2). https://doi.org/10.5751/ES-12306-260208

Sandbrook, C., Luque-Lora, R., & Adams, W. (2018). Human Bycatch: Conservation Surveillance and the Social Implications of Camera Traps. https://doi.org/10.17863/CAM.30452

Schneider, S., Taylor, G. W., Linquist, S., & Kremer, S. C. (2019). Past, present and future approaches using computer vision for animal re-identification from camera trap data. Methods in Ecology and Evolution, 10(4), 461–470. https://doi.org/10.1111/2041-210X.13133

Sethi, S. S., Ewers, R. M., Jones, N. S., Orme, C. D. L., & Picinali, L. (2018). Robust, real-time and autonomous monitoring of ecosystems with an open, low-cost, networked device. Methods in Ecology and Evolution, 9(12), 2383–2387. https://doi.org/10.1111/2041-210X.13089

Sturley, S., & Matalonga, S. (2020). PANDI: A Hybrid Open Source Edge-based System for Environmental and Real-Time Passive Acoustic Monitoring - Prototype Design and Development. 2020 1st International Conference on Innovative Research in Applied Science, Engineering and Technology (IRASET), 1–6. https://doi.org/10.1109/IRASET48871.2020.9092006

Swanson, A., Kosmala, M., Lintott, C., & Packer, C. (2016). A generalized approach for producing, quantifying, and validating citizen science data from wildlife images. Conservation Biology, 30(3), 520–531. https://doi.org/10.1111/cobi.12695

Swanson, A., Kosmala, M., Lintott, C., Simpson, R., Smith, A., & Packer, C. (2015). Snapshot Serengeti, high-frequency annotated camera trap images of 40 mammalian species in an African savanna. Scientific Data, 2(1), 150026. https://doi.org/10.1038/sdata.2015.26

Tabak, M. A., Norouzzadeh, M. S., Wolfson, D. W., Sweeney, S. J., Vercauteren, K. C., Snow, N. P., Halseth, J. M., Salvo, P. A. D., Lewis, J. S., White, M. D., Teton, B., Beasley, J. C., Schlichting, P. E., Boughton, R. K., Wight, B., Newkirk, E. S., Ivan, J. S., Odell, E. A., Brook, R. K., … Miller, R. S. (2019). Machine learning to classify animal species in camera trap images: Applications in ecology. Methods in Ecology and Evolution, 10(4), 585–590. https://doi.org/10.1111/2041-210X.13120

Terada, S. (2021). Building human–elephant relationships based on science and local ownership: A long-lasting issue in the era of Sustainable Development Goals. Animal Conservation, 24(5), 738–739. https://doi.org/10.1111/acv.12742

UN General Assembly. (2015). Transforming our world: The 2030 Agenda for Sustainable Development A/RES/70/1. https://www.refworld.org/docid/57b6e3e44.html

Walker, K. L. (2012). Labor costs and crop protection from wildlife predation: The case of elephants in Gabon. Agricultural Economics, 43(1), 61–73. https://doi.org/10.1111/j.1574-0862.2011.00565.x

Wearn, O. R., & Glover-Kapfer, P. (2019). Snap happy: Camera traps are an effective sampling tool when compared with alternative methods. Royal Society Open Science, 6(3), 181748. https://doi.org/10.1098/rsos.181748

Whytock, R. C., & Christie, J. (2017). Solo: An open source, customizable and inexpensive audio recorder for bioacoustic research. Methods in Ecology and Evolution, 8(3), 308–312. https://doi.org/10.1111/2041-210X.12678

Whytock, R. C., Świeżewski, J., Zwerts, J. A., Bara-Słupski, T., Pambo, A. F. K., Rogala, M., Bahaa-el-din, L., Boekee, K., Brittain, S., Cardoso, A. W., Henschel, P., Lehmann, D., Momboua, B., Opepa, C. K., Orbell, C., Pitman, R. T., Robinson, H. S., & Abernethy, K. A. (2021). Robust ecological analysis of camera trap data labelled by a machine learning model. Methods in Ecology and Evolution, 12(6). https://doi.org/10.1111/2041-210X.13576

